# Newly regenerated axons through a cell-containing biomaterial scaffold promote reorganization of spinal circuitry and restoration of motor functions with epidural electrical stimulation

**DOI:** 10.1101/2020.09.09.288100

**Authors:** Ahad M. Siddiqui, Riazul Islam, Carlos A. Cuellar, Jodi L. Silvernail, Bruce Knudsen, Dallece E. Curley, Tammy Strickland, Emilee Manske, Parita T Suwan, Timur Latypov, Nafis Akhmetov, Shuya Zhang, Priska Summer, Jarred J. Nesbitt, Bingkun K. Chen, Peter J. Grahn, Nicolas N. Madigan, Michael J. Yaszemski, Anthony J. Windebank, Igor Lavrov

**Author notes:** Corresponding author: Igor Lavrov, MD, PhD, Department of Neurology, Department of Physiology and Biomedical Engineering, Mayo Clinic, 200 First Street SW, Rochester, MN, 55905. Co-first author (Ahad M. Siddiqui, Riazul Islam).

## Abstract

We report the effect of newly regenerated neural fibers via bioengineered scaffold on reorganization of spinal circuitry and restoration of motor functions with electrical epidural stimulation (EES) after spinal transection (ST). Restoration across multiple modalities was evaluated for 7 weeks after ST with implanted scaffold seeded with Schwann cells, producing neurotrophic factors and with rapamycin microspheres. Gradual improvement in EES-facilitated stepping was observed in animals with scaffolds, although, no significant difference in stepping ability was found between groups without EES. Similar number of regenerated axons through the scaffolds was found in rats with and without EES-enabled training. Re-transection through the scaffold at week 6, reduced EES-enabled motor function, remaining higher compared to rats without scaffolds. The combination of scaffolds and EES-enabled training demonstrated synaptic changes below the injury. These findings indicate that sub-functional connectivity with regenerated across injury fibers can reorganize of sub-lesional circuitry, facilitating motor functions recovery with EES.

---

Current therapies for spinal cord injury (SCI) demonstrate only limited functional recovery. Recent advances in bioelectronics and tissue engineering make available novel neuromodulatory and neuroregenerative therapies, aimed to improve function after SCI^1-5^. Encouraging results using epidural electrical stimulation (EES) to restore motor function in humans with SCI^6-8^ have been attributed to the presence of functionally silent fibers and the combination of EES and motor training^9-11^. Most of the patients diagnosed with complete loss of motor function after SCI demonstrate some degree of sub-functional connectivity, which provides the anatomical and functional basis for supraspinal control in the presence of EES^11,12^. Elucidating the mechanisms of EES-enabled restoration of motor function after SCI with sub-functional connectivity is critical for optimizing combined neurostimulation and neuroregenerative therapy. In this study we report the first evidence of the contribution of axons that have regenerated through a scaffold in the reorganization of spinal circuitry and restoration of motor functions with EES.

The transplantation of scaffolds with GDNF secreting Schwann cells^13^ or with rapamycin and Schwann cells^14^ have shown functional and anatomical improvement after SCI. As an experimental platform for this study, we combined hydrogel scaffolds composed of positively charged oligo-[poly(ethylene glycol)fumarate] (OPF+) loaded with neurotrophic factor (GDNF) secreting Schwann cells and rapamycin microspheres (enhanced scaffolds) combined with EES and EES-enabled motor training (Fig. 1A-C)^15-19^. Groups with SCI and EES-facilitated training (*EES only*) (n=4), with enhanced scaffolds *(scaffold only)* (n=3), and with *combined therapy* of scaffold and EES-enabled training (n=10, 5 with re-transection) were compared (Fig. 1B).

**Figure 1:**
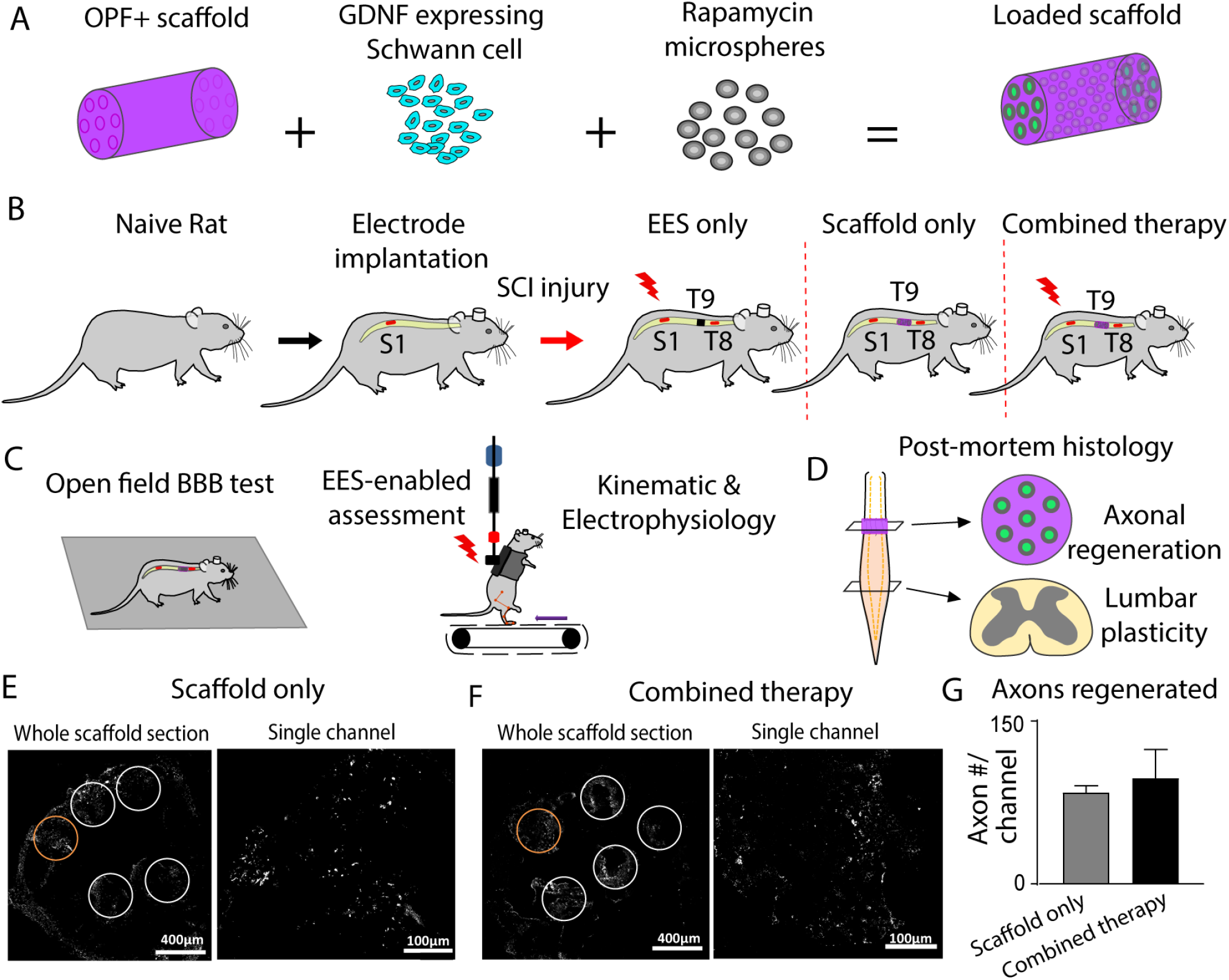
Experimental methods and study design. (A) Preparation of 7 channel OPF+ hydrogel scaffold loaded with GDNF expressing Schwann cell and rapamycin. (B) *In vivo* experimental model with implantation of the electrodes and scaffold following complete transection (T9). There were 3 groups of rats that received a T9 transection with epidural electrical stimulation (EES only), T9 transection with scaffold placement (scaffold only), and transection with scaffold placement and EES (combined therapy). (C) Following SCI and implantations, all rats received EES-enabled motor training on a treadmill with outcome collected as open field BBB score, kinematic, and electrophysiology during EES enabled stepping on treadmill. (D) At the end of the experiment, axonal regeneration through the scaffold and change in plasticity across lumbosacral spinal cord were assessed. (E) Transverse section (10 µm thick) through a scaffold implanted in a rat that did not receive EES and (F) transverse section in rat that received EES-enabled motor training. The sections were fixed and stained with β-tubulin 7 weeks post implantation and axons were visualized and quantified in the channels (white dotted circles). The channels in the orange dotted circles are presented in higher magnification on the right. (G) The number of axons passing through the scaffold shows no statistical difference between group without EES (n=3) and group that received EES-enabled motor training (n=5) (t-test).

At 7 weeks after complete SCI, the number of axons passing through each of the seven channels of the scaffold was assessed (Fig. 1D-G; Supplementary Fig 1). Groups with scaffold only (Fig. 1E) and with combined therapy (Fig. 1F) demonstrated regeneration through the scaffold, although, with no difference between the groups: 84.46±5.93 and 97.60±26.16, axons/channel respectively (Fig. 1G).

## Figure 1 about here

EES-enabled motor performance was evaluated with kinematics, EMG, and electrophysiology. Open field behavior *Basso, Beattie, and Bresnahan (BBB) motor scores* were collected with and without EES. BBB scores at week 1 without stimulation were similar across all groups (0-1.5) with no difference during the first 4 weeks. At week 4 and 6 the BBB score was greater for rats with combined therapy (wk4: 5.29±0.71; wk6: 6.62±1.38) compared to EES only (wk4: 2.62±0.63, p<0.05; wk6: 2.50±0.67, p<0.05). *Kinematic analysis* (Fig. 2A-E) at 2-4 weeks demonstrated higher step height in the group with combined therapy compared to animals with scaffold only (p<0.05) and with EES only (p<0.001). At week 6 the group with scaffolds only recovered step height almost to the level of group with combined therapy. Both groups with scaffold only and with combined therapy demonstrated higher step height compared to EES only (p<0.05) (Fig 2C). An increase in step length was found in the group with combined therapy at 2-4 weeks compared to scaffold only (wk2: p<0.05, wk4: p<0.01) or with EES only (wk2 and 4: p<0.01; Fig 2D). At week 6, the group with scaffolds only recovered to the level of the group with combined therapy. At 6 weeks after SCI, dragging was greater in the group with EES only compared to both combined therapy (p<0.001) and scaffold only (p<0.05) (Fig. 2A). All groups demonstrated recovery of angular displacement that in knee was higher at week 4 and 6 in group with combined therapy compared to other groups (p<0.05). MTP angles were not different at 2 weeks after injury, although at 4-6 week, the group with combined therapy demonstrated higher MTP displacement compared to other groups (p<0.05) (Fig. 2E).

**Figure 2:**
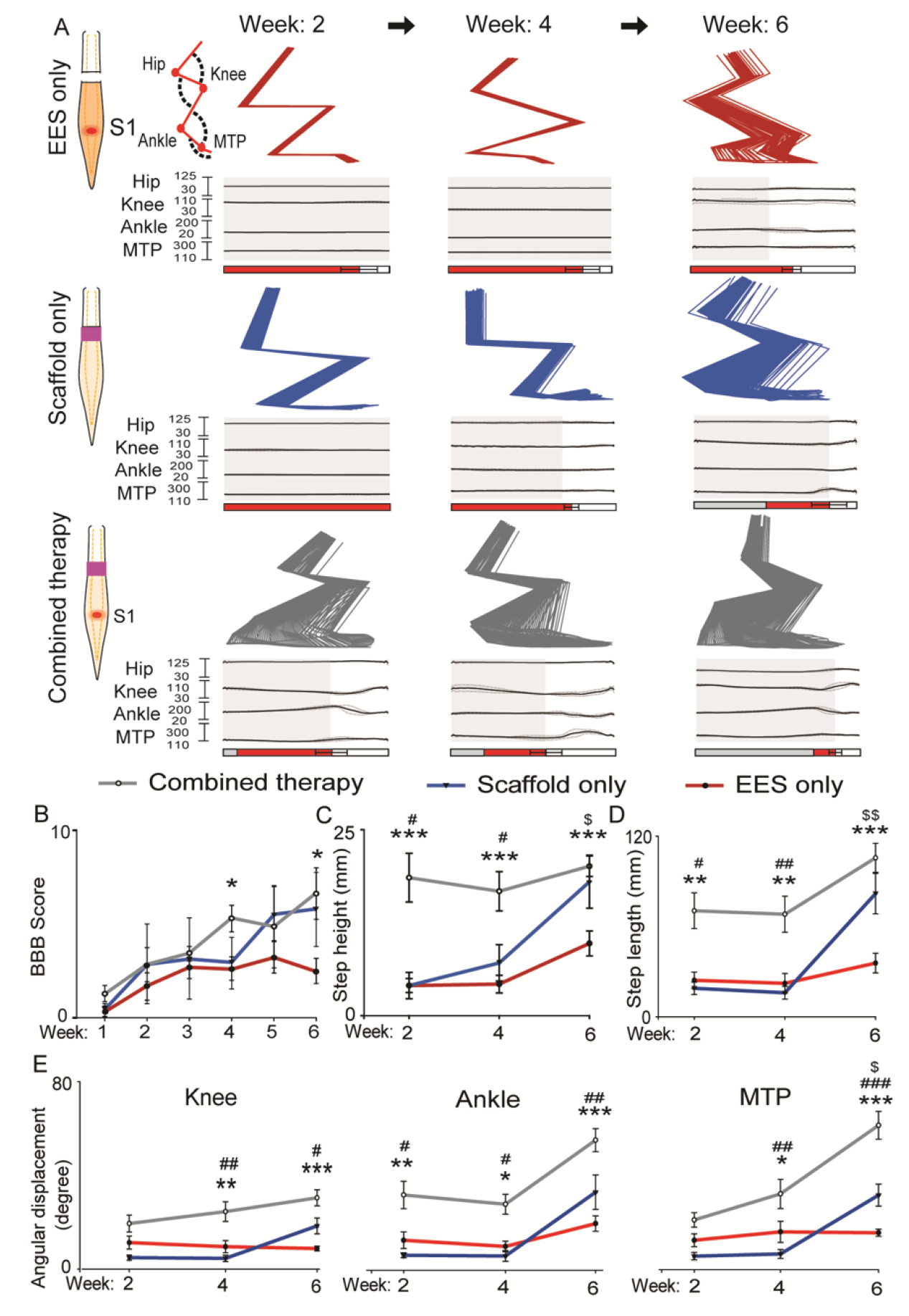
Rats receiving combined therapy showed early and sustained recovery. (A) Three representative examples of EES-enabled gait function recovery in animals after complete transection (T9) receiving EES only, Scaffold only or Combined therapy, collected at up to 6 weeks. Examples of stick diagram and joint angles (hip, knee, ankle, and MTP) averaged from 5 consecutive steps on treadmill. Stance phase is represented by gray rectangles, drag by red rectangles, and swing by white rectangles. (B) Motor performance assessed with BBB score in rats with EES only (n=4), with Scaffold only (n=3), and with Combined therapy (n=5) was accessed for 6 weeks post injury (*p <0.05, multiple t-test). Gait kinematic parameters, step height (C) and step length (D), were compared between the groups at 2, 4 and 6 weeks after spinal cord transection. At each time points these parameters were found to be significantly improved in rats with Combined therapy, compared to the rats with EES only. Displacement of joint angles, knee, ankle, and MTP (E) (*p <0.05, **p<0.01 and ***p<0.001, one-way ANOVA and post hoc analysis using Holm-Sidak method. Outcome presented EES only vs. Combined therapy^*^, with scaffold only vs. with combined therapy^#^, with EES only vs. with Scaffold only^$^).

## Figure 2 about here

*Spinal Cord Motor Evoked Potentials (SCMEP)*^*20*^ were recorded while rats were stimulated above (T8) and below (S1) the injury at week 2 and 6, and after re-transection at week 7 (Fig 3A). In a representative rat with combined therapy, stimulation above the injury (T8) at week 2 evoked small Supralesional Evoked Polysynaptic Response (SEPR) (Fig. 3A). By week 6 SEPR became more prominent and disappeared after re-transection. Stimulation below the injury at S1 evoked middle response (MR) (4.5-5.7ms) that increased in amplitude following re-transection, indicating the influence of regenerated fibers on the sub-lesional network. In another rat, with stimulation at T8, SEPR was detected only at week 6 and also disappeared after re-transection (Fig. 3A). Stimulation below the injury at S1 in this rat also demonstrated facilitation of MR response after re-transection. The mean MR peak-to-peak amplitude before re-transection at week-6 was 4.28 mV and increased to 10.67 mV after re-transection (p<0.001) (Fig 3B).

**Figure 3:**
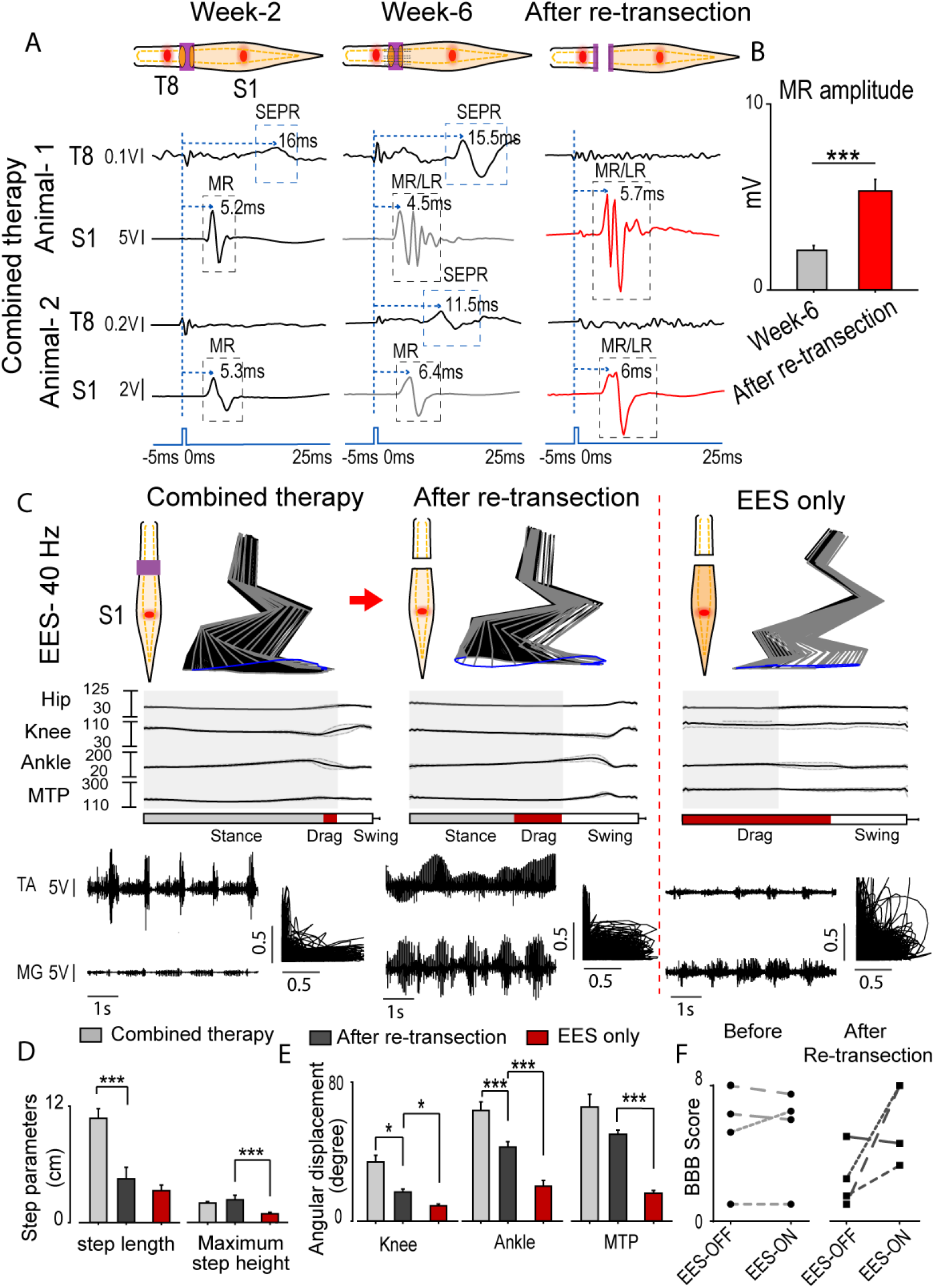
(A) Representative example of averaged Spinal Cord Motor Evoked Potential (SCMEP) (n=8 responses) of rats implanted with hydrogel scaffold, and received EES-enabled motor training, collected at 2 and 6 weeks after spinal transection and after re-transection across the scaffold following 6 weeks of recovery and potential regeneration through the scaffold. SCMEPs were recorded from hind-limb muscle medial gastrocnemius (MG), while spinal cord was stimulated using epidural electrode placed at T8 segment (above the injury) and at S1 segment (below the injury). Examples of SCMEPs were collected from representative animal-1 & animal-2. Blue dotted line indicates the moment when EES pulse was applied. Middle and late responses (MR/LR) and Supralesion Evoked Polysynaptic Response (SEPR) are indicated with grey and blue rectangles. (B) Peak-to-peak amplitude of MR at week-6 and after re-transection. (C) Animals with scaffolds and EES (Combined therapy) recovered greater angular displacements of the knee, ankle, and MTP. It is important to note, that after re-transection at week 6, improvement in motor function was greater compared to transected animals without scaffold (EES only). The stance phase is represented by grey rectangle, drag by red, and swing by white. (D) Comparison of gait parameters, step length and maximum step height. (E) Angular displacements of hip, knee, ankle, and MTP. (F) BBB score assessed with EES OFF and ON before and after re-transection (n=4).The same rats are tracked before and after re-transection using same types of dashed line. (Data are represented as mean +/- standard error. *p <0.05, **p<0.01 and ***p<0.001, one-way ANOVA and post hoc analysis using Holm-Sidak method).

## Figure 3 about here

*Electrophysiological and behavioral features* of motor performance were compared during EES-enabled motor training during 6 weeks after SCI and after re-transection. Rats with combined therapy demonstrated gradual recovery of EES-enabled stepping and after re-transection they retained some improvements compared to EES only group (Fig. 3C). Following re-transection, rats with combined therapy lost step length (p<0.001) (Fig 3D) as well as had decrease in knee and ankle angles, that still were greater compared to EES only group (p<0.05) (Fig 3E). The MTP angle did not change after re-transection, although, was greater (p<0.001) compared to EES only (Fig 3E). Interestingly, rats with combined therapy demonstrated similar BBB scores with EES turned OFF and ON, although, after re-transection, BBB score with EES-ON was increased for 3 of the 4 rats compared to when the EES was OFF.

*Morphological reorganization* below the injury was evaluated with synaptophysin co-localized to ChAT positive motor neurons and calbindin positive interneurons (spinal segments L2-S1) (Fig. 4A-D). Rats with combined therapy (0.46±0.088) had greater co-localization of ChAT and synaptophysin compared to EES only group (0.28±0.073; p<0.05; Fig. 4E). Greater synaptophysin co-localized to calbindin was found in rats with combined therapy (0.45±0.07) compared to EES only (0.31±0.062; p<0.05; Fig. 4F). When the distribution of synaptophysin vesicles was quantified on ChAT+ neuronal cell bodies, the EES only group had significantly lower numbers (38,615 synaptophysin vesicles per ChAT+ neuron/mm^2^, mean: 54,486,CI_L_: 50,993 CI_U_: 53,776) compared to the animals from the combined therapy group (78,656, mean: 93,287, CI_L_: 89,534 CI_U_: 97,039; p<0.0001) and with scaffold only (53,387, mean: 66,105, CI_L_: 60,893 CI_U_: 71,317; p<0.001). The amount of synaptophysin vesicles on ChAT+ neuronal cell bodies was greater in rats with combined therapy compared to scaffold only (p<0.0001; Fig. 4G). Similar difference was found in the distribution of synaptophysin vesicles on calbindin+ cell bodies (Fig 4H). The amount of synaptophysin was greater in rats with combined therapy (118,512 synaptophysin vesicles per calbindin+ neuron/mm^2^, mean: 127,661, CI_L_: 122,482 CI_U_: 132,839; p<0.0001) compared to EES only (59,914, mean: 66,562, CI_L_: 62,201 CI_U_: 70,924). The amount of synpatophysin vesicles in rats with scaffold only (77,738, mean: 92,142, CI_L_: 85,732 CI_U_: 98,551) was intermediate to the EES only (p<0.0001) and combined therapy groups (p<0.0001). These results indicate that the regenerating axons affect synaptic reorganization below SCI, although, the combined therapy has greater influence on reorganization on the motor and interneuron cells bodies. The synaptophysin distribution was greater on calbindin+ interneurons than the ChAT+ motor neurons (p<0.0001), suggesting the greater role of synaptic reorganization on interneurons with the ratio of synaptophysin per neuron/mm^2^ on ChAT+ to calbindin+ greater for the scaffold only (0.68) and combined therapy (0.66) than the EES only group (0.59).

**Figure 4:**
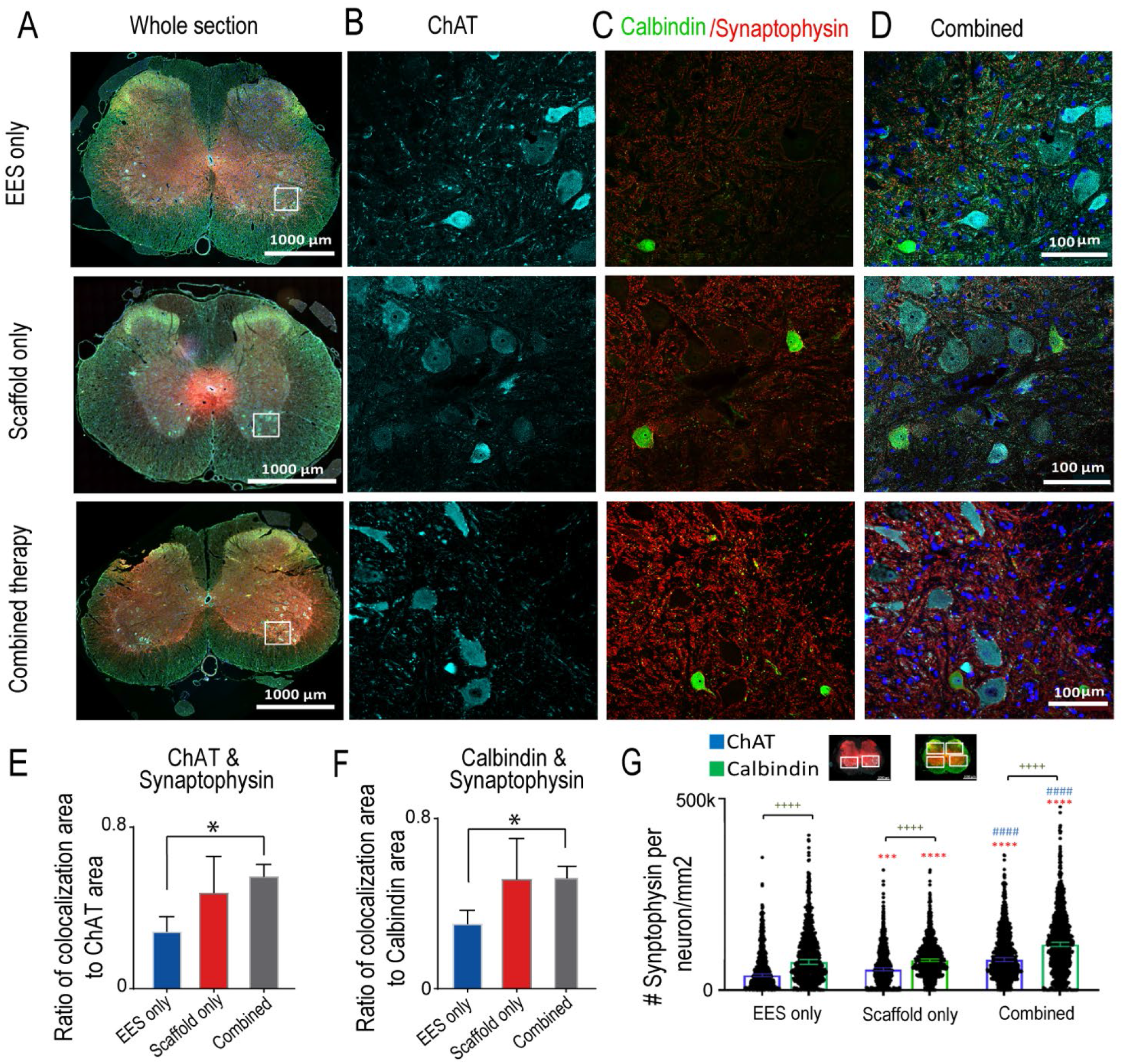
Morphological reorganization of spinal cord below the injury was observed in rats receiving combined therapy. Changes in synaptic vesicle co-localization with motor neurons and interneurons in the lumbosacral spinal cord of rats with EES only, with Scaffold only and with Combined therapy. (A) Image of a transverse section (10 µm thick) of the whole spinal cord taken at 20x and tiled between spinal segments L2-S1. The spinal cords were immunostained for, (B) choline acetyltransferase (ChAT) positive motor neurons(cyan), (C) calbindin positive interneurons (green) and synaptophysin (red). (D) Combined image with all immunostains and DAPI nuclear staining (blue). (E) Average area of ChAT co-localized with synaptophysin in rats with EES only (n=3), with Scaffold only EES (n=3), and with Combined therpay (n=5). (F) Similar comparison for Calbindin co-localized with synaptophysin (t-test, EES only vs. Combined therapy, and Scaffold only vs. Combined therapy). (G) Distribution of synaptophysin on ChAT+ motor neuron (EES only: n=913, Scaffold only: n=812, combined therapy n=1086 neurons) and Calbindin+ interneuron (EES only: n=420, Scaffold only: n=954, combined therapy n=1315 neurons) cell bodies per cell area. The bars are shown as median +/- confidence interval. Statistical comparisons were made using Kruskal-Wallis test with Dunn’s multiple comparison (* compared to EES only, # compared to Scaffold Only, + comparison between ChAT and Calbindin; * p<0.05, ** p<0.01, *** p<0.001, **** p<0.0001).

## Figure 4 about here

These results demonstrate the effect of newly regenerated axons on motor performance facilitated with EES through synaptic reorganization of sub-lesional circuitry, suggesting that neuroregenerative and neuromodulatory therapies cumulatively improve motor function after SCI. Reconnection across complete SCI has been achieved with a nerve autograft and scaffolds^21,22^, particularly when delivering cell types and growth factors^22-28^. Although the functional outcome in this study was better in rats who received combined therapy, the number of axons through scaffolds was not different between the groups and was in between previously reported with scaffold loaded GDNF secreting Schwann cells^25^ or rapamycin with Schwann cells^29^. Rapamycin may reduce the number of regenerating axons, although previous study demonstrated improvement when compared with OPF+ scaffolds loaded with Schwann cells alone^14^. This could be attributed to altering synaptic excitability^30-33^. Rapamycin can block mTOR required for axon regeneration^34,35^, although its impact on behavior is not evident^36-39^. BBB score at 6 weeks of combined therapy in this study (6.62±1.38) was greater compare to OPF+ with rapamycin microspheres and Schwann cells (4.69±0.57)^14^. 4 week time-point in this study of the combined therapy group (5.29±0.7) was similar to effect of OPF+ scaffolds with GDNF Schwann cells (3.67±0.4)^13^. Limitations of BBB^40,41^ we added to the validity of these findings with electrophysiological and kinematic analysis in behavior assessment system^42^. Improvement in group with combined therapy vs. EES only, and with intermediate effect in scaffolds only group was found. Following re-transection, group with combined therapy still demonstrated better performance compared to EES only group. The difference between EES only vs. combined therapy group indicates on the role of newly regenerating axons in reorganization of sub-lesional circuitry. Greater synaptophysin co-localization with motor neuron and interneuron in animals treated with combined therapy suggests that regenerating axons form functional connections with sub-lesional circuitry facilitating reorganization with EES-enabled training. Plasticity below the SCI has been attributed to rehabilitation^43,44^, while the quality of movement is largerly depends on descending commands and sensory feedback integration on different interneuronal populations^11,17,45-48^. Although the numbers of rats are relatively small, the concordance of behavioral, electrophysiological and histological results convincingly demonstrate that greater synaptic reorganization on the interneurons in the presence of regenerated axons leads to improvements in polysynaptic responses, improving gait, particularly in group with combined treatment. In summary, the results of this study for the first time demonstrate that newly regenerating through the cell-containing scaffold axons with EES-enabled motor training can reorganize sub-lesional circuitry and improve EES-enabled motor performance, providing a platform for synergistic testing and translation of regenerative and neuromodulatory therapies to maximize functional restoration after SCI.

## Methods

Seventeen Adult female rats (Sprague–Dawley, 250–300 g body weight) were used in this study. All rats were implanted with stimulating electrodes epidurally at T8 and S1 spinal cord levels and EMG electrodes implanted in the hind limb muscles tibialis anterior (TA) and medial gastrocnemius (MG). The groups studied include, (1) rats implanted with EES electrodes but no Scaffold and received EES (EES only; n=4), (2) rats implanted with EES electrodes and scaffold but received no EES (scaffold only; n=3) and (3) rats implanted with EES electrodes and scaffold and received EES-enabled motor training (combined therapy; n=10). In order to study the functional effect of fibers regenerated across the scaffold, group-3 animals were retransected 6 weeks after complete SCI at level T9.

### Surgical procedures

The rats were deeply anesthetized by a combination of ketamine (100 mg/kg) and Xylazine (10 mg/kg) administered intraperitoneally (IP) and maintained at a surgical level with supplemental doses of ketamine as needed. Buprenorphine was administered as a single dose at the beginning of the experiment. All surgeries were performed under aseptic conditions. All procedures involving animals were approved by the Mayo Clinic Institutional Animal Care and Use Committee and all guidelines were followed in accordance with the National Institute of Health as well as Institute for Laboratory Animal Research and the United States Public Health Services Policy on the Humane Care and Use of Laboratory Animals.

### Electromyography wire and electrode implants for SCI rats

A small skin incision was made at the midline of the skull. The muscles and fascia were retracted laterally, and the skull was thoroughly dried. A 12-pin Omnetics circular connector (Omnetics, Minneapolis, MN) and 12 Teflon-coated stainless-steel wires (AS632, Cooner Wire, CA) were attached to the skull with screws and dental cement as previously described ^20,49,50^. Skin and fascia incisions were made to expose the bellies of the MG and TA muscles bilaterally. Using hemostats, the EMG wires were routed subcutaneously from the back incision to the appropriate locations in the hind-limb. Bipolar intramuscular EMG electrodes were inserted into the muscles as described previously ^20,50^. The EMG wires were coiled near each implant site to provide stress relief. Electrical stimulation through the head-plug was used to visually verify the proper response of the electrodes in each muscle ^20^. A partial laminectomy was then performed at the L2 vertebral level (S1 spinal segment) and one wire was affixed to the dura at the midline using 9.0 sutures as previously described ^49^. A small notch made in the Teflon coating (about 0.5–1.0 mm) of the wire used for EES was placed toward the spinal cord and served as the stimulating electrode. Another laminectomy was performed to expose the area for implantation of the electrode at T8 spinal segment. The wire was coiled in the back region to provide stress relief. Teflon coating (about 1 cm) was stripped from the distal centimeter of one wire that was inserted subcutaneously in the back region and served as a common ground ^20,51^.

### Poly-lactic-co-glycolic acid (PLGA)-rapamycin microsphere fabrication

Microspheres fabricated from PLGA were used to slowly elute the antifibrotic drug rapamycin. The polymer used to form the PLGA microspheres was 50:50 lactic acid to glycolic acid with 29 kDA molecular weight (Resomer RG 503 H; Sigma-Aldrich). An oil-in-water emulsion with solvent evaporation technique as previous described ^14,52,53^. Briefly, 1 mg of rapamycin was dissolved in 100 µL of absolute ethanol. The rapamycin-ethanol solution vortex emulsified dropwise for 30 seconds in 250 mg PLGA dissolved in 1 mL of methylene chloride. The mixture was then emulsified in 2 mL of 2% (w/v) poly vinyl alcohol for 30 seconds. This solution was then mixed with 100 mL of 0.3% (w/v) poly vinyl alcohol and 100 mL of 2% (w/v) isopropanol and stirred for 1 hour to evaporate methylene chloride. The microspheres are then centrifuged at 2000 rpm for 3 minutes and washed 3 times with distilled water and centrifugation. The liquid is then discarded and the microspheres are frozen at −80 °C for 1 hour. Lastly, the microspheres are dried overnight under vacuum and then used for scaffold manufacture.

### Scaffold Preparation

Positively charged OPF+ scaffolds with seven channels (Fig. 1A) were fabricated as previously described ^14,54,55^. Briefly, the liquid polymer consist of 1g of OPF macromere (16,246g/mol) dissolved in 650 µL of deionized water, 0.05% (w/w) photoinitiator (Irgacure 2959; Ciba Specialty Chemicals) 0.3 g N-vinyl pyrrolidinone (NVP; Sigma), and 20% [2-(methacryloyloxy) ethyl]- trimethylammonium chloride (MAETAC; Sigma). To embed the rapamycin microspheres, 25 mg of microspheres was stirred into 250 µL of OPF+ polymer liquid. The liquid polymer containing the microspheres is then mold injected over 7 equally spaced wires with 290 µm diameter in a glass cylinder. After 1 hour exposure the UV light (365 nm) ay 8 mW/cm^2^, the individual scaffolds are cut into 2mm lengths. Before use, they are serially washed in ethanol 3 times for 30 minutes each followed by 4 1X PBS washes ^56^.

### Primary and GDNF-secreting Schwann cells

Glial derived neurotrophic factor (GDNF) secreting SCs were created as described previously ^25^. Briefly, primary rat SCs were harvested from the sciatic nerve of Sprague Dawley pups between postnatal day 2 and 5. The nerves are stripped of connective tissue and the epineurium and cut into 1mm sections. Then then are enzymatically treated with 0.25% trypsin EDTA (Mediatech Inc.) and 0.03% collagenase (Sigma). Then the cells are mechanically dissociated by pipetting and centrifuges for 5 minutes at 188 G. The primary SCs were then genetically modified by seeding 80,000 cells/mL and grown with high titer (2×1010 cfu) retroviral (GDNF-eGFP) supernatant media with 8 µg/mL polybrene for 24 hours. Transduced SCs were selected for 12 days with 1 mg/ml G418 analogue in SC media containing 50:50 DMEM: F12 media (GIBCO) supplemented with 10% FBS, 1% antibiotic-antimycotic, 2µM Froskilin (Sigma), and 10 ng/mL neuregulin-1 (recombinant human NRG-1; R&D Systems)) at 37^°^C in 5% CO_2_. The cells were expanded and frozen until ready for use.

### Spinal cord transection and scaffold implantation

One week after electrode implantation surgery, a complete spinal cord transection was performed. Rats were anaesthetized with a mixture of oxygen and Isoflurane (≈1.5%). Mid-dorsal skin incision was made between T6 and L4 and the paravertebral muscles were retracted as needed. A partial laminectomy was performed at the T8 level and the dura was opened longitudinally. Lidocaine was applied locally and the spinal cord was completely transected using a micro-scissors. Completeness of the lesion was verified by two surgeons under microscope. Then, some rats were implanted with GDNF/SC-RAPA-OPF+. Buprenorphine (0.5–1.0 mg/kg subcutaneous injections twice a day) were administered for analgesia. Tissues were sutured by layers and animals were allowed to recover in individual cages with soft bedding. Manual bladder expression was performed four times daily during two weeks post transection. The rats were allowed to move over an open field surface to help stimulate their bladder and reflexes once daily for 2 weeks. The hind limbs of the spinal rats were also moved passively through a full range of motion once per day to maintain joint mobility.

### Spinal cord electrical epidural stimulation

A single channel manually controllable isolated stimulator (A-M systems, Sequim, WA) or an eight independent channel real-time programmable (STG4008, Multichannel Systems, Reutlingen) stimulator were used to deliver biphasic square wave electrical stimulation (250 μs pulse width) at 40 Hz with amplitudes ranging from 0.5 to 2.5 V to the epidural electrode placed on the rat’s lumbosacral (S1) spinal cord to facilitate motor activity.

### Training and animal care

Rats were acclimated to a specially designed motor-driven rodent treadmill and body weight support system ^42^ for a period of 7 days prior to surgery for 15-20 minutes each day (Fig. 1B). One week post-surgery, the rats went through a manual bipedal step training rehabilitation process (30 min a day, 3 days a week) for 6 weeks under the influence of EES at 40 Hz subthreshold level (0.4 to 2 V). Chronic step training was used because it helps to engage and reinforce the locomotor networks.

### BBB assessment

The Basso, Bettie, and Bresnahan (BBB) locomotor rating scale was used to access the hind-limb function weekly starting one week post-injury (Fig. 1C). Three independent observers were blinded to the animal groups and score was given on the 21 point scale. The BBB scoring was performed as previously described ^57^, except that the animals were scored twice with and without subthreshold EES (40Hz). At least 30 minutes of rest was given between the sessions. The BBB scores from the independent observers were averaged and then the left and right limb scores were averaged for each rat. Multiple t-test was used to determine statistical difference between groups.

### Kinematics

A motion tracking system (Vicon, UK) was used to record three-dimensional digital position of back and the hind limb joints at 100 Hz. Six motion-sensitive Infra-Red (IR) cameras were aimed at the treadmill or the open field volumes. Another high-speed video camera synchronized with motion tracking system was positioned in a side to provide a lateral view of the motor performance. Retro-reflective markers were placed on bony landmarks at the iliac crest, greater trochanter, lateral condyle of the femur, lateral malleolus and the distal end of the fifth metatarsal on both legs of the rat to record the kinematics of the hip, knee and ankle joints. Nexus system was used to obtain three-dimensional coordinates of the markers. Analysis of kinematic data was performed using method previously described ^42^.

### Electrophysiology

EMG activity was collected from TA and MG muscles on both legs at 4000 Hz during stepping and later high-pass filtered at 0.5 Hz to remove direct current offset. In order to determine flexor (TA) and extensor (MG) coordination, EMG signals were band pass filtered (10 Hz to 1000 Hz), rectified, normalized and plotted on X and Y axis consecutively ^42^. The excitability of the spinal cord were determined by stimulating the spinal cord at S1 spinal level while *spinal cord motor evoked potentials (SCMEP)* were recorded from the hind-limb muscles via EMG^20^, while the rats were suspended in the body weight support system ^42^. The functional connectivity across the injury and reflexes below the injury were tested by stimulating T8 spinal level above and S1 segment below the injury, while recording SCMEP from hindlimb muscles. During testing, the rats were suspended allowing hind limbs to freely respond and stimuli of increasing intensity (amplitude 0.5-4.5 V, pulse width 0.25 ms, frequency 0.5 Hz) were applied to epidural spinal electrodes. 10 consecutive responses were collected during spinal cord stimulation above (T8) and below (S1) the injury with a single pulse to evaluate SCMEP in awake animals to assess the influence of information delivered across scaffold on SCMEP during standing and during stepping on a treadmill.

### Perfusion and dissection

Seven weeks post injury that rats were transcardially perfused using 4% paraformaldehyde (PFA) in phosphate buffered saline (PBS). The spinal column was removed en block and post-fixed in 4% PFA in PBS for 2 days at 4°C. The spinal columns were then washed 3 times in PBS and dissected out. Then 6 mm segment containing the scaffold at the center and adjacent spinal cord was embedded in paraffin. A 1 cm block covering L2-S1 spinal cord segment was also embedded in paraffin. Transverse or longitudinal serial sections of 10 μm thick were made on a Reichert-Jung 23 Biocut microtome (Leica, Bannockburn, IL) of the scaffold and spinal cord areas.

### Immunohistochemistry

Post-mortem immunohistochemistry was performed in all groups of animals in order to assess regeneration through the scaffold and plasticity change in lumbosacral spinal cord (Fig. 1D). Primary antibodies used were against β-III tubulin (Tuj-1; mouse anti-rat, 1:300; Millipore), choline acetyltransferase (ChAT; goat anti-rat. 1:50, Millipore), Calbindin (CB38; rabbit, 1:1000, SWANT), and synaptophysin (mouse, 1:50, Abcam). The secondary antibodies used were against mouse Cy3 (donkey, 1:200, Jackson ImmunoResearch Laboratories), goat Alexa 647 (donkey, 1:200, Jackson ImmunoResearch Laboratories), rabbit Cy2 (donkey, 1:200, Jackson ImmunoResearch Laboratories). The slides were deparaffinized through serial washes through xylene, ethanol, and distilled water. Antigen retrieval was performed through incubation in 1 mM EDTA in PBS for 30 minutes in a rice steamer. The sections were then blocked using 10% normal donkey serum in 0.3% Triton X-100 in PBS for 1 hour. Next, TrueBlack (Biotium) was used as per manufacturer’s instructions to quench autofluorescence due to lipofuscin. Then, the sections were incubated overnight in the primary antibody diluted in 5% normal donkey serum in PBS at 4° C. The next day after 3 washes in PBS, the sections were incubated in secondary antibody diluted in PBS containing 5% normal donkey serum for 1 hour at room temperature. Lastly, the sections were washed 3 times in PBS containing, and then mounted with SlowFade Gold Antifade Reagent with DAPI (Molecular Probes, Eugene, Oregon, USA).

### Axon Count

Axons were identified as punctate staining of β-III tubulin and were counted in transverse paraffin sections at quarter lengths throughout the scaffold. Tiled 20x images of the whole scaffold were imaged using a Zeiss LSM 780 inverted confocal microscope. StereoInvestigator software suite (MBF) was used to count the axons in each channel using Optical Fraction Probe. Briefly, a square grid measuring 100µm x 100µm was overlaid on the image after contouring around the channel. The axon was counted if it fell either in the quadrant frame or overlapped with the acceptance lines (top horizontal and right vertical boundary). The average axon counts per channel for each quarter length were averaged yielding one value per animal. The total axon counts of rats with scaffolds and with/without EES were compared using two sample t-test.

### Colocalization of synaptophysin with neuronal populations

Antibody against ChAT was used to identify motor neuron populations, while Calbindin was used to identify interneuron subtype on sections taken from transverse and longitudinal spinal cords between spinal segments L2-S1. The co-localization tool in Image-Pro (Media Cybernetics) was used to determine the amount of overlap of synaptophysin with ChAT or Calbindin according to manufacturer’s instructions. Color pairs were created between the red (synaptophysin) and green (Calbindin) channels and the red and far red channels (ChAT). A threshold was applied using the auto-bright feature. The process was standardized and automated to have consistent, unbiased analysis. The analysis was done using 40x images taken on a Zeiss LSM 780 inverted confocal microscope serially through the lumbosacral spinal cord every 1500 µm. On each slide 2 sections 20 µm apart were chosen and on each section 4 areas were chosen and titled: the left and right dorsal horn and ventral horn (Supplementary Fig. 2A). The areas of interest were then averaged per section, the 2 sections per slide were averaged, and then all the slides were averaged to yield one average per animal. The amount of co-localization was compared between animals with scaffolds and EES that were re-transected and no scaffold and EES using an unpaired t-test.

Distribution of synaptophysin on ChAT+ and calbindin+ cell bodies was analyzed using a semiautomated image analysis system on the QuPATH open source software ^58^ with the images described above. The cell bodies labelled with ChAT or calbindin were first traced (Supplementary Fig. 2B). The cell area of the traced space was recorded. Then the cell detection tool was used to identify the synaptophysin vesicles. The detection channel was set for the red channel and then optimized parameters were used (requested pixel size 0 µm, background radius 0 µm, median filter radius 1 µm, sigma 0.5 µm, minimal area 0.01 µm^2^, maximum area 5 um^2^, threshold 30). The number of synaptophysin vesicles per neuron were normalized to cell body area and all the neurons were plotted. Then the median with upper 95% confidence interval (CI_U_) and lower 95% confidence interval (CI_L_) was used to indicate the peak of the distribution. Kruskal-Wallis test with Dunn’s multiple comparison was used to analyze the distribution.

### Statistical analysis

*S*tatistical analysis was performed using SigmaPlot (Systat Software, UK) or Prism (GraphPad). The data were first tested for normality and then one-way Analysis of Variance (ANOVA) was performed to determine if the groups had different outcomes. If the groups were found significantly different (P< 0.05) a post-hoc analysis was performed using Holm-Sidak method to compare against control and p values were reported. When compared between two groups, two samples t-test was performed. All results were reported as mean ± standard error, and ^*, #, $^P< 0.05, ^**, ##, $$^P< 0.01 and ^***, ###, $$$^P< 0.001, unless otherwise noted. Non parametric analysis was done using Kruskal-Wallis test with Dunn’s multiple comparison.

## Funding sources

Minnesota State Office for Higher Education Spinal Cord Injury and Traumatic Brain Injury Research Grant Program (FP00093993), North American Spine Society, Morton Cure Paralysis Fund, and the Bowen Foundation.

## Figures and Legends

**Supplementary Figure 1:**
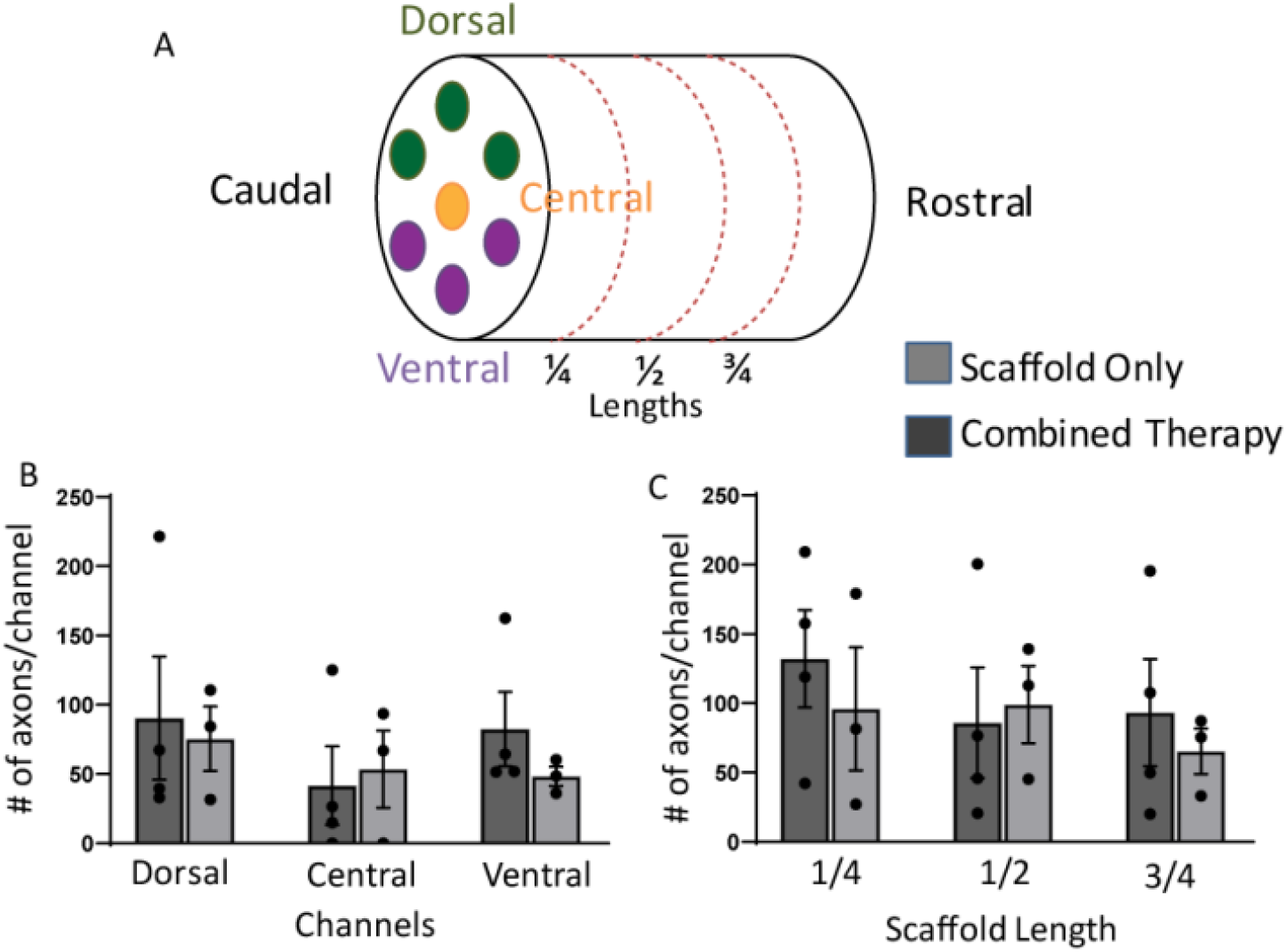
(A) Scaffold sectioning method applied in the experiment. Each scaffold was sectioned at ¼, ½ and ¾ lengths and the number of axons at each channel were counted in all three sections. The three top channels on the dorsal side are identified as dorsal (green), the central channel (orange) and the three channels on the ventral side is identified as ventral (purple). (B) Number of axons regenerated per channel in the dorsal, central, and ventral channels are compared between the rats with scaffold only and rats receiving combined therapy. (C) Similar comparison of number of axons regenerated per channel at ¼, ½ and ¾ lengths of the scaffold.

**Supplementary Figure 2:**
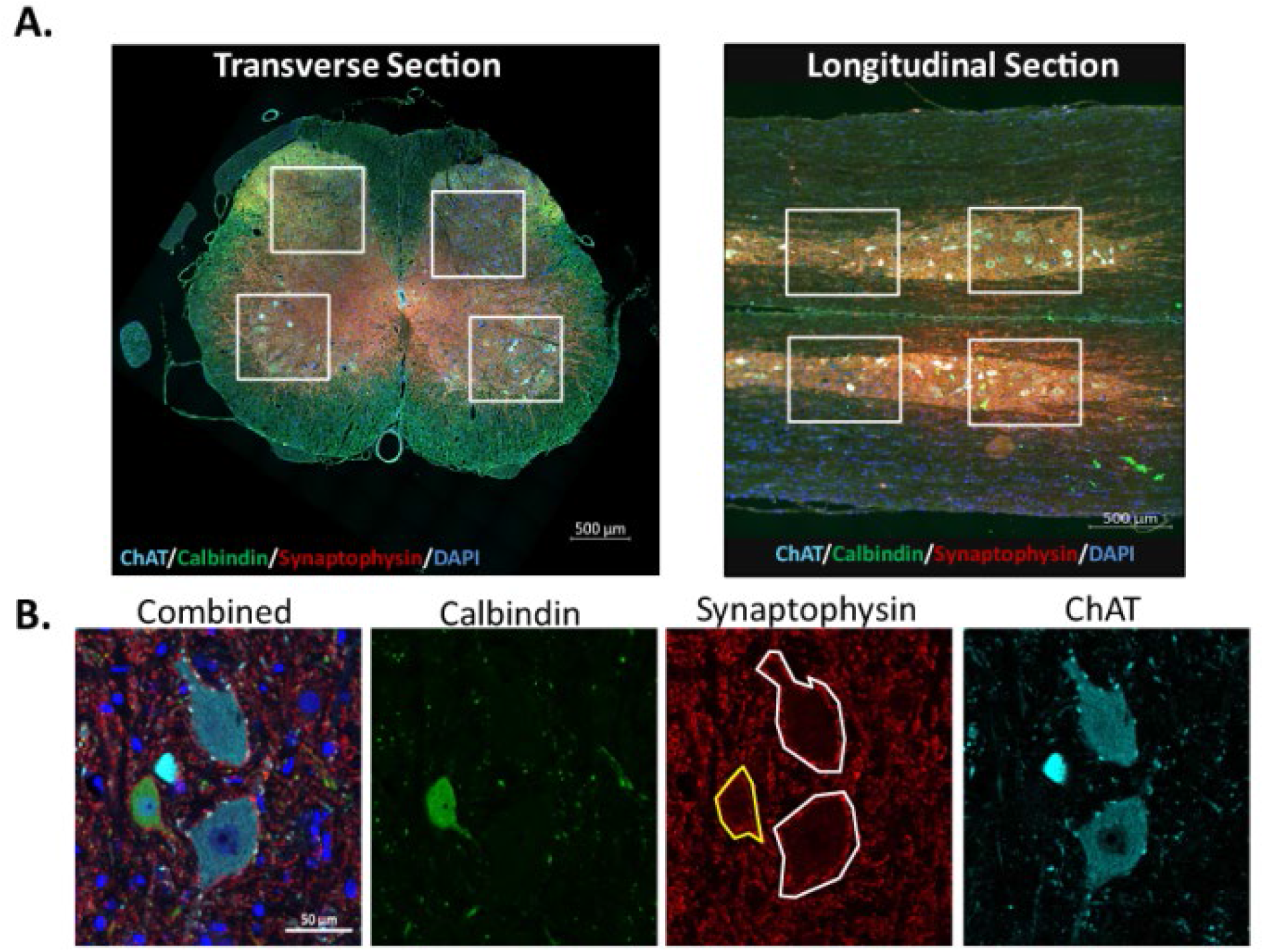
(A) Example of transverse and longitudinal sectioning of lumbosacral spinal cord. The spinal cord sections are co-stained with ChAT, Calbindin, Synpatophysin and DAPI. Four regions (enclosed by white squares) in the dorsal and ventral regions were selected to perform analysis on plasticity change due to regeneration through the scaffold. (B) Representative example of immunohistochemistry performed on the lumbosacral spinal cord, combined DAPI, Calbindin, Synaptophysin, ChAT and individual stained images shown at 40x zoom. The area around the cell body were traced (white tracing for ChAT positive cells and yellow for calbindin positive cells) to analyze the amount of synaptophysin in the cell boundary.

## Notes

### Competing Interest Statement

The authors have declared no competing interest.

